# Statistical analysis of no observed effect concentrations or levels in eco-toxicological assays with overdispersed count endpoints

**DOI:** 10.1101/2020.01.15.907881

**Authors:** Ludwig A. Hothorn, Felix M. Kluxen

## Abstract

In (eco-)toxicological hazard characterization, the No Observed Adverse Effect Concentration or Level (NOAEC or NOAEL) approach is used and often required despite of its known limitations. For count data, statistical testing can be challenging, due to several confounding factors, such as zero inflation, low observation numbers, variance heterogeneity, over- or under-dispersion when applying the Poisson model or hierarchical experimental designs. As several tests are available for count data, we selected sixteen tests suitable for overdispersed counts and compared them in a simulation study. We assessed their performance considering data sets containing mixing distribution and over-dispersion with different observation numbers. It shows that there is no uniformly best approach because the assumed data conditions and assumptions are very different. However, the Dunnett-type procedure based on most likely transformation can be recommended, because of its size and power behavior, which is relatively better over most data conditions as compared to the available alternative test methods, and because it allows flexible modeling and effect sizes can be estimated by confidence intervals. Related R-code is provided for real data examples.

## 1 Introduction

While the use of statistical tests have been recently criticized [2], [36], many (eco-)toxicological assays require the derivation of no observed adverse effect concentrations or levels (NOAEC, NOAEL which are used interchangeable in the current document) by statistical test methods. Sometimes the primary endpoints are count data, such a number of offspring in daphnia aquatic assays, with several replicates per concentration [34], which are not normal-distributed. The variability between these replications should be modeled accordingly, e.g. as a variance component in the mixed effect model or as an over-dispersed discrete distribution.

Recently the use the closure principle computational approach test (CPCAT) assuming the Poisson distribution was proposed [23] for the analysis of count data, e.g. reproduction data in (eco-) toxicology. While a simulation study on power was conducted in order to verify its statistical power [22], the test was not compared to other test methods and only considered limited assumption violations. This is concerning, because the analysis of count data can be challenging, due to several confounding factors:

- A possible zero inflation, e.g. in the control group of a micronucleus assay with near-to-zero counts
- Possible over- or even under-dispersion compared to the standard Poisson model
- An additional variance component between the experimental units (plates, flasks, tanks)
- Possible variance heterogeneity between the concentrations
- Very different value ranges of the counts (either very small values in direction to ordered categorical values including zero, or large values in direction of approximate normal distribution)

In practice, these effects often occur jointly. All of those confounding factors affect the reliability of derived p-values or confidence intervals if not considered in the applied statistical test method, which can result in unreliable hazard characterization estimates.

Further, for count data in assays with small observations numbers (*n*_*i*_, denoted sample size in biostatistics) a distribution assumption can neither be determined by pre-tests (because of too small sample sizes *n*_*i*_) nor by mechanistic arguments of the underlying biological process (due to missing relationship between model and distribution type). Therefore and in general, a flexible and robust approach to specific assumptions or data situations would be helpful, instead of a test that is only exactly optimal for one selected assumption. As several tests are available for count data, we selected 16 tests suitable for overdispersed counts and compared them in a simulation study. Here the focus is on modifications of the Dunnett test [4] for the comparison of several treatments to a control group that provides effect size and their simultaneous confidence intervals whenever possible. We assessed their performance considering data sets containing mixing distribution and over-dispersion with different observation numbers.

It shows that there is no uniformly best approach because the assumed data conditions and assumptions are too different. The most likely transformation method [17, 14] can be recommended from the point of view of flexible modeling and robustness. And, because effect sizes can be easily derived, which is generally recommended [15]. We further describe two challenges in more detail when comparing count data, which relate to data generation and the definition of the NOAEC/L.

## 2 Motivating examples

Three motivating examples with count endpoints were selected and presented by boxplots also contain the raw data [27]. In the left panel, the number of offsprings in a daphnia bioassay [1], in the middle panel the number of micronuclei in a micronucleus assay on phenylethanol [5], and in the right panel the number of revertants in a TA98 Salmonella Ames assay [25] are presented. We see i) counts in a small value range or in a larger one, ii) variance heterogeneity (both increasing or almost zero), iii) approximately symmetrical distributions in designs up to 5 doses and iv) small *n*_*i*_ down to even 3.

Not just global overdispersion (i.e. variance higher than Poisson variance) occur in these data, but dose-specific dispersions and/or variance heterogeneity between the concentrations.

**Figure 1:**
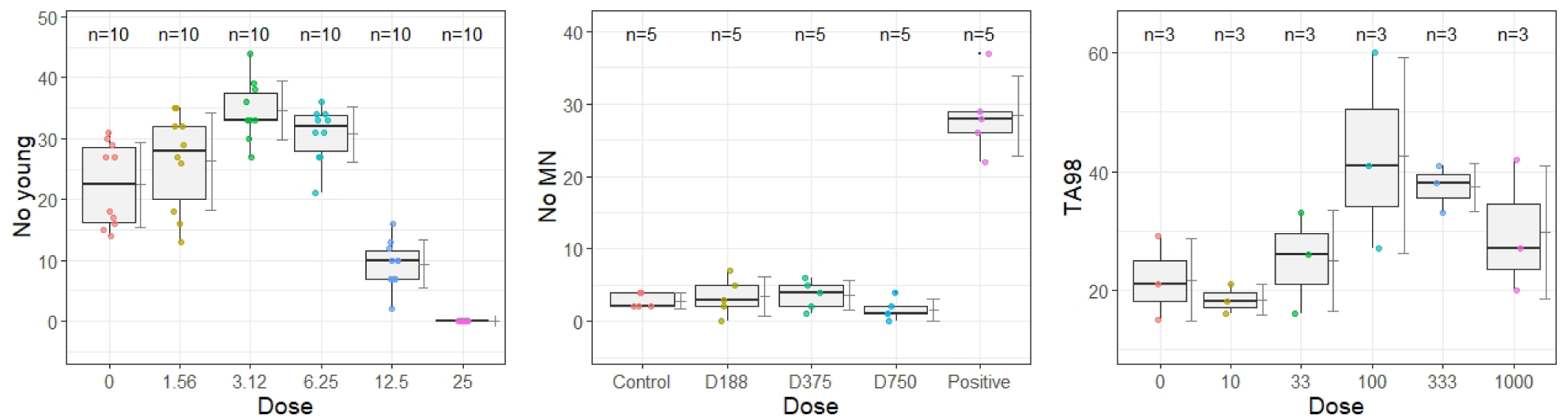
Box plots with superimposed individual values corresponding means with standard deviations of three example data sets from literature. Left) Shows the number of offspring (No young) from a daphnia bioassay [1]. Middle) Shows the number of micronuclei (No MN) of a genotoxicity assay conducted with phenylethanol [5]. Right) Shows the number of revertants (TA98) in on strain of an Ames assay [25].

## 3 Approaches and initial considerations

A problem is the unclear definition of the NOAEC in a randomized design, which usually includes several concentrations *NC*(*C*_0_ = 0), *C*_1_, …, *C*_*k*_ (where k commonly 3-5). The actually direct method is the estimation of the maximum safe dose 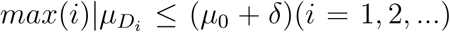 [13, 32, 33], i.e., by directed non-inferiority tests. This is not used in (eco-)toxicology at all, probably because a consensus for endpoint-specific non-inferiority threshold *δ* is lacking, i.e. an agreed effect size which is not considered toxicologically relevant. However, also the assumption of NOAEC via *NOAEC* = *D*_*MED*_ − 1 is problematic because of the direct control of the false positive error rate, and also because the definition of the minimum effective dose (MED) is blurred: significant or relevant decision, assuming a concentration-response monotonicity into account or not, using hypothesis tests or nonlinear models, modeling concentration qualitatively [9] or quantitatively [29, 3].

### 3.1 NOAEC definition

Two definitions are used here:

A. Just the concentration before the lowest significant one

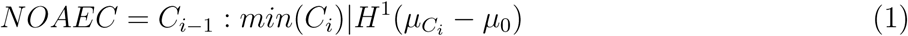

where all higher dose effects are ignored.
B. The concentration before the lowest significant concentration where all higher ones must be significant as well-but not necessarily monotonous ordered.

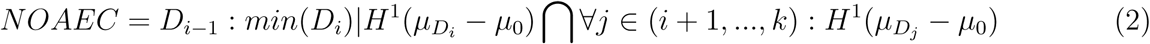

Both definitions can be extended for effective concentrations, i.e. 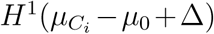 which is commonly not used because of the missing fixation of Δ. One-sided tests (or more appropriate confidence limits) are throughout where the direction of harm is a-priori known.

Note, contrary to the both definitions, the no observed adverse effect level is also defined as *the highest dose tested that causes no adverse effects in a test species in a properly designed and executed study* [6], which becomes relevant when no statistically observed effect levels exist between statistically observed effect levels.

### 3.2 NOAEC procedures

Following eq(1) definition, the following four hypotheses tests can be performed: i) Closed testing procedure (CTP) [10], ii) Dunnett-type procedure [12], iii) step-down pairwise tests [7], and iv) contrast tests [28].

CTP for eq(1) can be formulated for comparison against control only, forming a complete family of hypotheses [31], which does no need complex partition-type hypotheses. For *k* = 3 and simple tree alternative the hypotheses are: global [0, 1, 2, 3], partition: [0, 1, 2], [0, 2, 3], [0, 1, 3], elementary [0, 1], [0, 2], [0, 3]; closed under intersection. While for monotonic order restriction the hypotheses are much simpler: [0, 1, 2, 3], partition: [0, 1, 2], elementary [0, 1] (because the rejection of [0, 1, 2, 3] implies the rejection of [0, 2, 3], [0, 1, 3], etc. under monotonic alternative. Any level *α*-test can be used for each hypothesis (including different tests for different hypotheses). Reflecting the many-to-one (NC) comparisons, the Helmert contrast seems to be suitable for simple tree alternatives, where under total order restriction two-sample tests *D*_*max*_ − *D*_0_ (or better related pairwise contrasts when the common degree of freedom is used) are suitable. ANOVA-type tests can be used as well, because of their inherently 2-sided hypotheses formulation they are not too appropriate [22]. A hybrid method robust for both monotone and non-monotone shapes is available as well [38].

Notice, model selection approaches [21] or Bayesian approaches [26] are not considered here, because they follow different testing philosophies, which would not allow an appropriate comparative assessment.

### 3.3 Possible biased estimations

At least six sources of biased NOAEC estimations exists, e.g. where the estimated NOAEC of one group is affected by the responses in other groups: i) NOAEC depends directly on *σ/n*_*i*_, both are no standardized (not so much in regulatory assays with at least *n*_*i*_ recommendation and a common variance), ii) NOAEC is an experimental concentration only (no model-specific interpolation), and depends on an appropriate choice of concentrations (number, intervals), iii) NOAEC based on a directional decision (a-priori clear either increase or decrease)-where two-sided testing may cause bias, iv) pooling contrasts/tests [8], [13], v) ignoring concentration-specific variance heterogeneity when using common MSE estimator, particularly in unbalanced designs, and vi) ignoring concentration-specific dispersions where common approaches focusing on a global dispersion parameter only. Figure 2 shows two common scenarios for such a bias. In the left panel *NOAEC* = *D*_1_, but the Williams contrast test [37] finds *MED* = *D*1 because of the significant pooled (*µ*_1_ + *µ*_2_ + *µ*_3_)*/*3 contrast. In the right panel *NOAEC* = *D*_1_, but the standard Dunnett test revealed *MED* = *D*_3_ because the extreme variance in *D*_3_ increases the common variance estimator and hence *D*_2_ is biased not significant. The related R-code is available in the Appendix.

**Figure 2:**
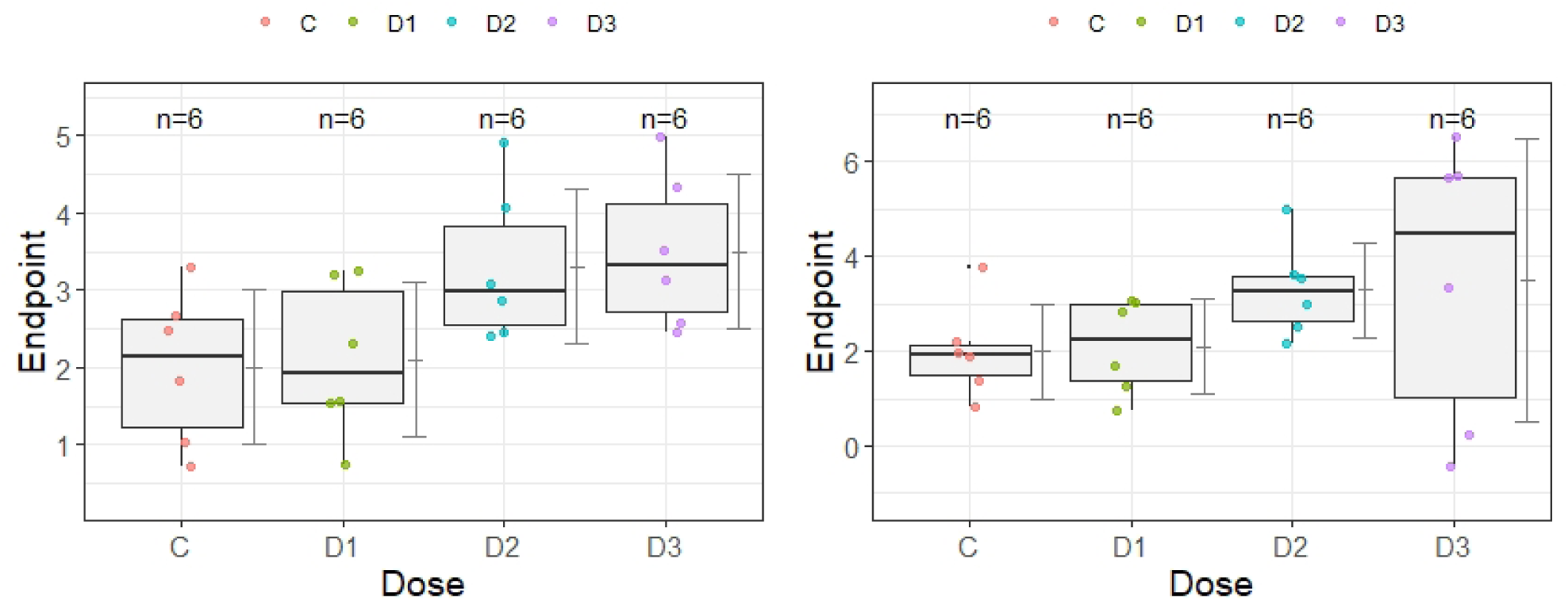
Box plots with superimposed individual values corresponding means with standard deviations of two simulated experiments with biased NOAEC estimation. The left panel shows a plateau-type shape where the Williams test estimates a too small NOAEC due to its pooling contrast issue. The right panel shows an increased variance in the high dose group resulting in a too large NOAEC estimation

## 4 Procedures comparing overdispersed counts in a design [*NC, C*_1_, …, *C*_*k*_]

As the standard approach, modifications of the Dunnett test [4] for comparing treatments versus a negative control in the generalized linear mixed effect model (glmm) [16] are commonly used. To model the variation between the experimental units, a quasi-Poisson model can be assumed or a random factor *between units* can be used in the mixed effect model. For the latter different algorithms exist, whereby the selection was made from the point of view of numerical stability at small *n*_*i*_. Alternatively a negative binomial model, not belonging to the glm family, can be assumed. Notice, these asymptotic approaches may be problematic for small *n*_*i*_ for both numerical stability and controlling the coverage probability. Therefore, a simple Freeman-Tukey transformation was used for the Dunnett test [11], [19] particularly appropriate for small *n*_*i*_. When the distribution is unknown and not only location effects should be considered, the most likely transformation model [12] can be used for the Dunnett test [14], the similar approach of continuous outcome logistic regression model [24] and the more specifically the cotram approach for count endpoints [30], i.e. most likely transformation methods for counts. A completely different approach is represented by the nonparametric Dunnett-type procedure for relative effect size, allowing any discrete distribution with variance heterogeneity [20]. The CPCAT test, a closed testing principle for parametric bootstrap tests [22] can also be conducted and has been proposed before for the assessment of (eco-)toxicological bioassays.

To account for variance heterogeneity between the concentrations, a sandwich variance estimator can be used when available, for linear models [39] as well as for generalized mixed effect models [35].

We compared 16 statistical test candidates in a simulation study, that cover the described different methods to analyse count data. Specifically the following tests were used:

1. Standard Dunnett test assuming normal distributed homoscedastic errors (Du)
2. Dunnett test modified with a sandwich variance estimator (DuS)
3. Dunnett test for log-transformed counts (DuL)
4. Dunnett test for log-transformed counts and a sandwich variance estimator (DuLS)
5. Dunnett test for Freeman-Tukey transformed counts (FT)
6. Dunnett test for Freeman-Tukey transformed counts and a sandwich variance estimator (DuFS)
7. Dunnett test in glm using Poisson link function (without overdispersion) (DuP)
8. Dunnett test in glm using quasi-Poisson link function (QP)
9. Dunnett test in the negative binomial model (DuN)
10. Dunnett test in the mixed effect model (MM)
11. Dunnett test in the mixed effect model using related sandwich estimator (DuMS)
12. Dunnett test for most likely transformation (MLT)
13. Dunnett test for continuous outcome logistic regression model (CORL)
14. Dunnett test for mlt-cotram (COT)
15. Dunnett test for continuous outcome logistic regression model (COTd)
16. Closure principle computational approach test (CPCAT)

In the appendix, the simulation results of these tests are presented in detail. After an initial study comparing 16 possible test candidates, we compared the most promising and relevant tests in a thorough simulation study, which is detailed in the following.

## 5 Simulation study

### 5.1 Size and any-pair power for comparisons vs. control

It is not clear from the start how to generate random count data in the (k+1) sample design with possible overdispersion and/or variance heterogeneity in the high(er) concentrations to characterize the size/power behavior of (eco-)toxicological assays. A first model assumes a mixing distribution of responding and non-responding subjects in the higher concentrations, based on integered normal mixing distributions (MIX). A second model based on mean-parametrized Conway-Maxwell-Poisson distribution allowing concentration specific dispersions [18] (CMP) For a fair comparison to CPCAT, 2-sided tests were considered only and tests with p-value outcome only.

#### 5.1.1 Results

Figure 3 shows two typical simulated scenarios. In the left panel an assay based on integered mixing distribution (*responder* + *non* − *responder*) (MIX) with overdispersion in the highest concentration is shown and in the right panel mean-parametrized Conway-Maxwell-Poisson distribution (CMP) in an inhibition assay with underdispersion in the highest concentration is displayed. Both balanced and unbalanced [*NC, C*_1_, *C*_2_, *C*_3_] designs were considered.

**Figure 3:**
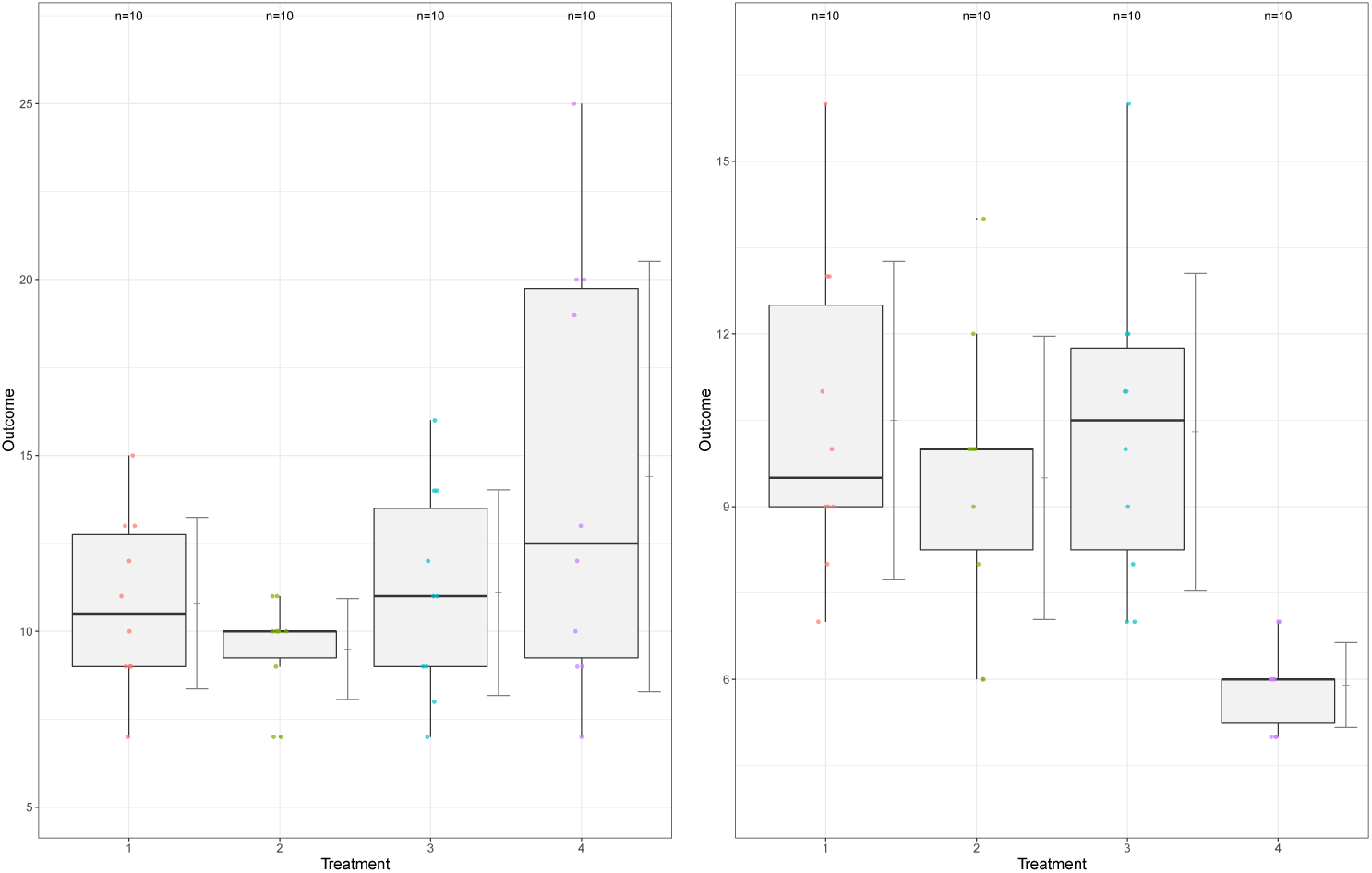
Example data sets in the simulation study under specific alternatives. The left panel shows a mixing distribution with overdispersion in the highest concentration. The right panel shows a mean-parametrized Conway-Maxwell-Poisson distribution with underdispersion in the highest concentration.

In the simulation study, the empirical size and any-pairs power (i.e. at least one contrast, anyone, in the alternative) were estimated from 16 test variants under different balanced and unbalanced designs (with medium and very small *n*_*i*_) for both distributions (MIX, CPM) with and without over-dispersion.

Appendix 1 shows the size (under the null hypothesis *H*_0_) and power (under certain alternative *H*_1_) estimates for various conditions for these 16 tests. Not surprisingly, there is no uniformly best test for all conditions. Table 1 summarized these estimated for the five preferred tests under different conditions: the Dunnett-tests for Freeman-Tukey transformed counts (FT), GLM with quasipoisson link function (QP), GLMM with Poisson link function (MM), most likely transformation (MLT) and as well the closed testing approach (CPCAT). Both site and power are not too different, i.e. from this point of view these 5 tests are all usable, The power functions for overdispersed CPM distribution in Figure 4 reveals a quite similar pattern under this special balanced *n*_*i*_ = 10 design, where MM and QP is more powerful than CPCAT over a wide range of non-centrality (effect size increase against control in a fixed design with constant variance).

**Table 1:** Size (under H0) and power (under H1) estimations for selected tests: (FT) Dunnett-tests for Freeman-Tukey transformed counts, (QP) generalized linear model with quasipoisson link function, (MM) generalized mixed effect linear model with Poisson link function, (MLT) most likely transformation, (CPCAT) the closed testing approach, under mixing distribution (MIX) or Conway-Maxwell-Poisson distribution (CMP) with and without overdispersion

| Distribution | Hypothesis | Dispersion | n | FT | QP | MM | MLT | CPCAT |
| --- | --- | --- | --- | --- | --- | --- | --- | --- |
| MIX | H0 | - | 10 | 0.06 | 0.01 | 0.07 | 0.08 | 0.001 |
|  | H0 | over | 10 | 0.06 | 0.07 | 0.02 | 0.06 | 0.01 |
|  | H1 | - | 10 | 0.91 | 0.87 | 0.40 | 0.88 | 0.16 |
|  | H1 | over | 10 | 0.50 | 0.60 | 0.38 | 0.51 | 0.22 |
| CMP | H0 | - | 10 | 0.05 | 0.06 | 0.04 | 0.07 | 0.04 |
|  | H0 | - | 5 | 0.06 | 0.08 | 0.04 | 0.11 | 0.03 |
|  | H0 | over | 10 | 0.05 | 0.05 | 0.05 | 0.06 | 0.05 |
|  | H0 | over | 5 | 0.06 | 0.10 | 0.07 | 0.11 | 0.08 |
|  | H1 | - | 10 | 0.89 | 0.91 | 0.90 | 0.91 | 0.85 |
|  | H1 | over | 10 | 0.84 | 0.87 | 0.88 | 0.85 | 0.84 |

**Figure 4:**
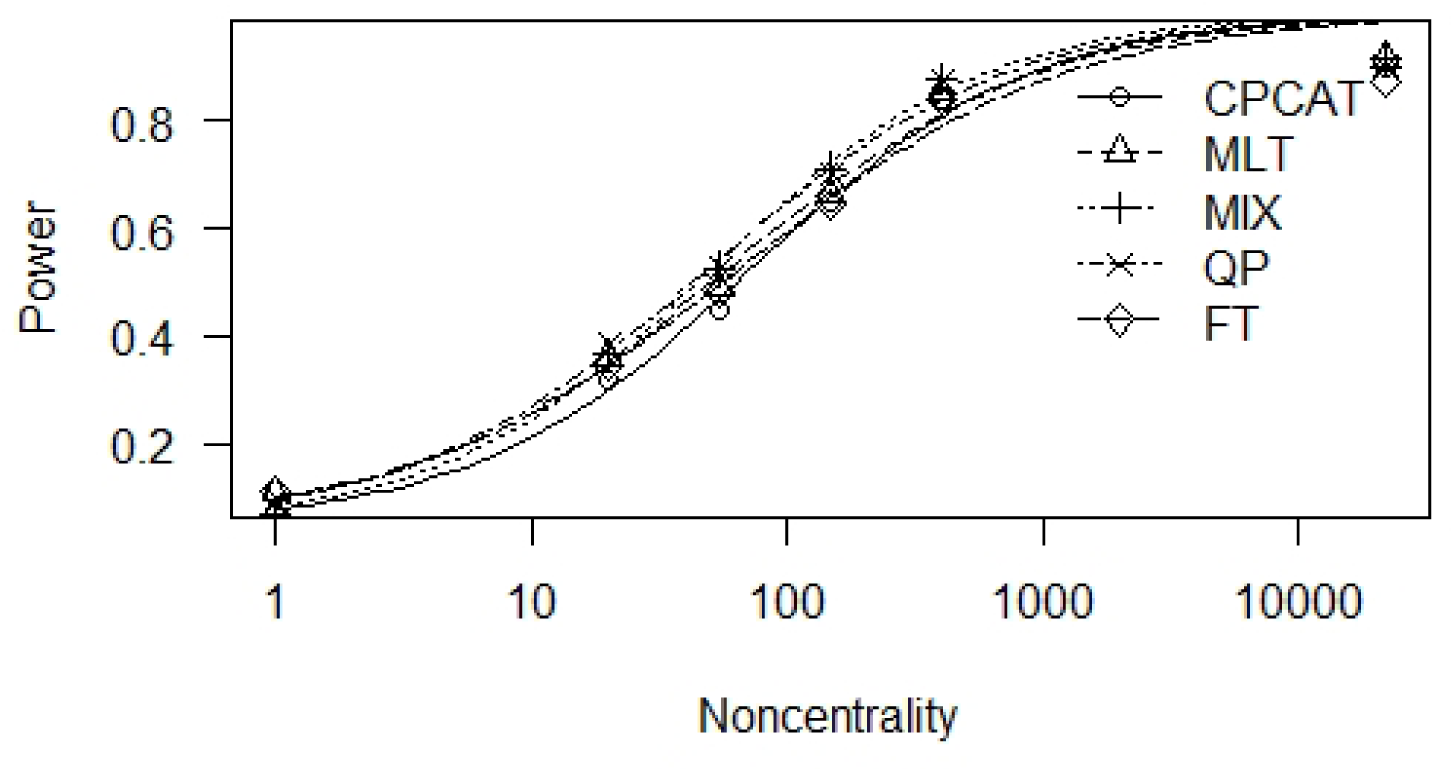
Power curve for selected tests: (FT) Dunnett-tests for Freeman-Tukey transformed counts, (QP) generalized linear model with quasipoisson link function, (MM) generalized mixed effect linear model with Poisson link function, (MLT) most likely transformation, (CPCAT) the closed testing approach. Power is (1-false negative error rate). Non-centrality is effect size increase against control

Notice, CPCAT is formulated for two-sided hypotheses only, whereas the other approaches can be formulated two-sided, or more appropriate, one-sided for NOAEC estimation. The impact on power is serious, e.g. for a certain point in *H*_1_ (CPM, overdispersed) the power of the two-sided MLT and CPCAT is 0.83, whereas the power increases to 0.90 for one-sided MLT Dunnett-type test.

### 5.2 Correct NOAEC estimation rates

The correct NOAEC rates for the above selected procedures (FT, QP, MM, MLT, CPCAT) were estimated for relevant concentration-response patterns (for 1000 runs and CPM distribution with overdispersion. The correct estimation rates are presented for selected conditions in Appendix II. Figure 5 shows a mosaicplot with the correct NOAEC estimation rates for decreasing non-centralities, abbreviated by C1, C2,…, C7. For condition C1 the true NOAEC is number 2 estimate, for condition C7 the true NOEAC is number 3 estimate where C2-C6 represent decreasing non-centrality in between. The patterns of the five tests are quite similar, with advantages for QP, MM and MLT.

**Figure 5:**
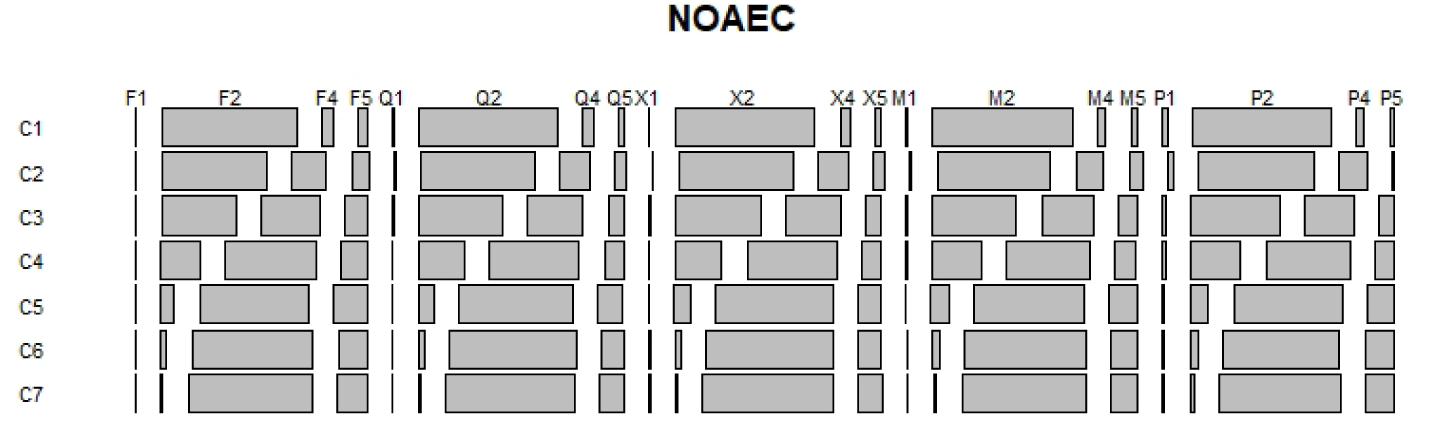
Mosaicplot for correct NOAEC estimation rates for 7 decreasing non-centralities C1,…,C7 for the selected tests FT, QP, MM, MLT, CPCAT.

## 6 Evaluation of two examples

The daphnia example was analysed with the five selected methods, where decreasing number of young is the relevant direction of harm. The related R-code is given in Appendix II. Table 2 reveals test statistics (t) and multiplicity-adjusted p-value for the five comparisons of the dose groups against control for the five selected tests (FT, QP, MM, MLT, CPCAT). While for FT, MLT and CPCAT the NOAEC is correctly estimated at 6.25, QP and MM show biased estimates due to variance heterogeneity and over-dispersion in the data.

**Table 2:** Test statistics (t) and adjusted p-value (p) of five selected tests FT, QP, MM, MLT, CPCAT of the daphnia example for the individual comparisons of dose against control

| Comparison | FT(t) | FT(p) | QP(t) | QP(p) | MM(t) | MM(p) | MLT(t) | MLT(p) | CPCAT(p) |
| --- | --- | --- | --- | --- | --- | --- | --- | --- | --- |
| 1.56-0 | 1.16 | 0.98 | 1.58 | 0.99 | 1.77 | 0.99 | -1.38 | 0.99 | 0.0818 |
| 3.12-0 | 4.69 | 0.99 | 4.53 | 0.99 | 5.07 | 0.99 | -4.21 | 0.99 | 0.0001 |
| 6.25-0 | 3.30 | 0.99 | 3.20 | 0.99 | 3.59 | 0.99 | -2.75 | 0.99 | 0.0013 |
| 12.5-0 | -5.51 | < 0.001 | -6.31 | < 0.001 | -7.07 | < 0.001 | 4.69 | < 0.001 | < 0.001 |
| 25-0 | -19.25 | < 0.001 | -0.01 | 0.89 | -0.04 | 0.89 | 6.72 | < 0.001 | < 0.001 |

The better representation are simultaneous confidence intervals, e.g. for the MLT approach (adapted to continuous outcome logistic regression modeling) where odds ratio are used as effect size, as demonstrated for the micronucleus assay example without positive control in Figure 6.

**Figure 6:**
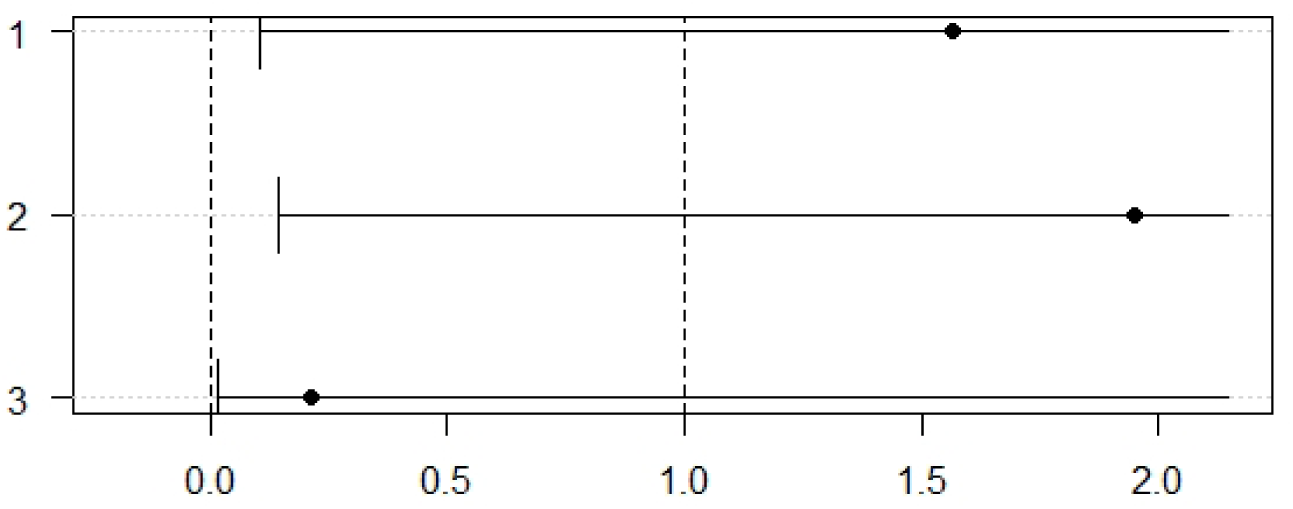
Odds ratios and their simultaneous confidence limits for the micronuceus example estimated by a MLT approach using continuous outcome logistic regression modeling

The dots representing the point estimators of the odds ratios (OR). The dotted line at 1 is for the null hypothesis of *OR* = 1. None of the lower limits is larger than 1, i.e. neither increase of number of MN in any dose can be concluded.

## 7 Conclusions

NOAEC/L estimation of count data in (eco-)toxicity assays poses a challenge, due to usually very small observation numbers, with overdispersed counts and an a priori unclear distribution assumption as well as possible variance heterogeneity between concentrations. We demonstrated that there is no uniformly best approach simply because the underlying data conditions and assumptions are very different between study designs. The most likely transformation methods can be recommended from the point of view of flexible modeling and robustness. They perform well in most data conditions. Procedures providing simultaneous confidence intervals, especially their interpretation as compatibility intervals, should be preferred to p values, as they allow the assessment of effect sizes which is recommended by several authors, e.g. [2, 15]. Our results also show that caution is advised when assays with too small *n*_*i*_, e.g. *n*_*i*_ = 3 are to be evaluated, which is similar to our previous observation for continuous data sets [14].

Corresponding R-code is included in the Appendix III for the examples and can be easily adapted to other data situations.

## 8 Appendix I: Detailed simulation results

Abbreviated: MIX …integered normal distributions, H0 … null hypothesis, H1… alternative hypothesis, d … using finite df, s… using sandwich estimator.

**Table 3:**
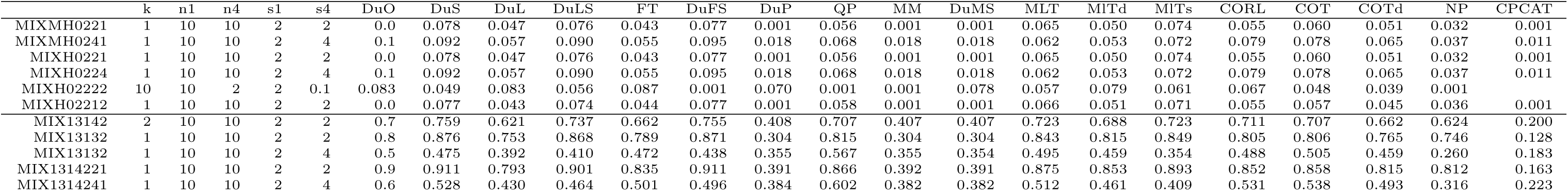
Simulated size (under H0) and power (under H1) for overdispersed integered mixing normal distributions

**Table 4:**
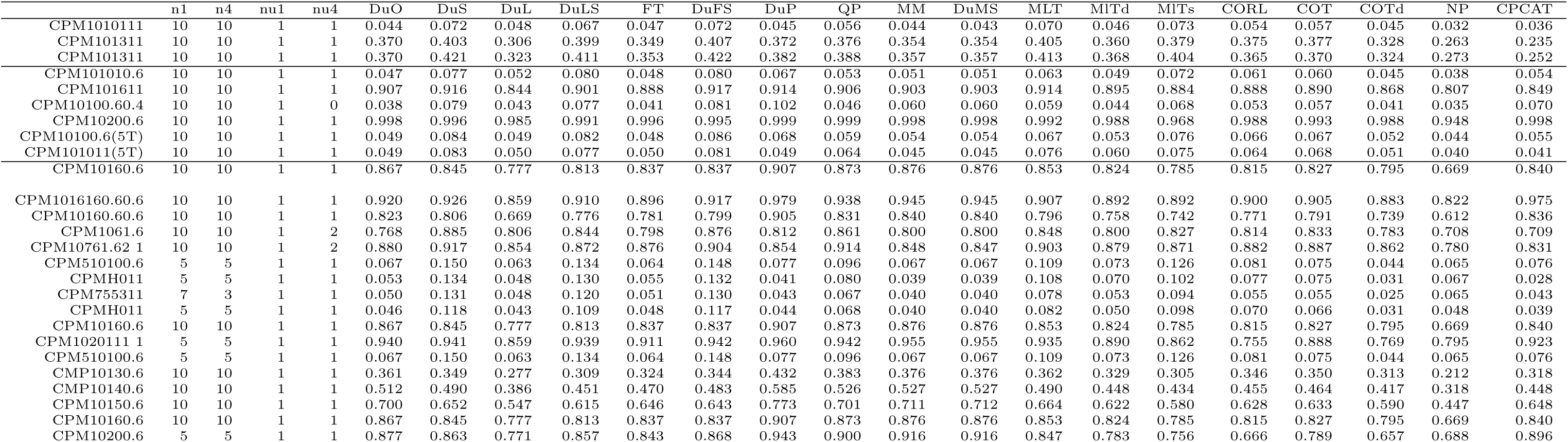
Simulated size (under H0) and power (under H1) for overdispersed CPM distribution

## 9 Appendix II: Correct NOAEC estimation rates

**Table 5:**
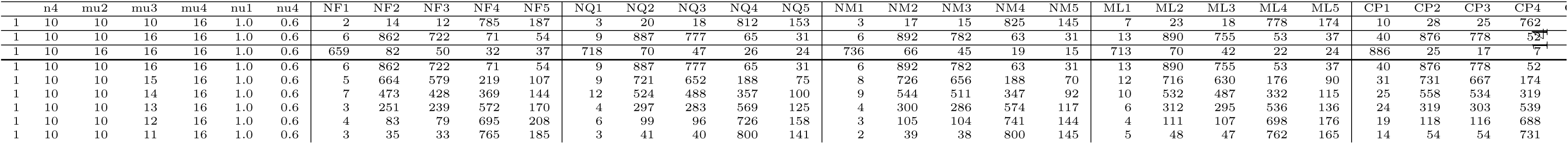
Correct NOEAC estimation rates (*NF*_*j*_ for FT-test, *NQ*_*j*_ for QP-test, *NM*_*j*_ for MM test, *ML*_*j*_ for MLT-test, *CP*_*j*_ for CPCAT-test

## 10 Appendix III: R-code of selected tests

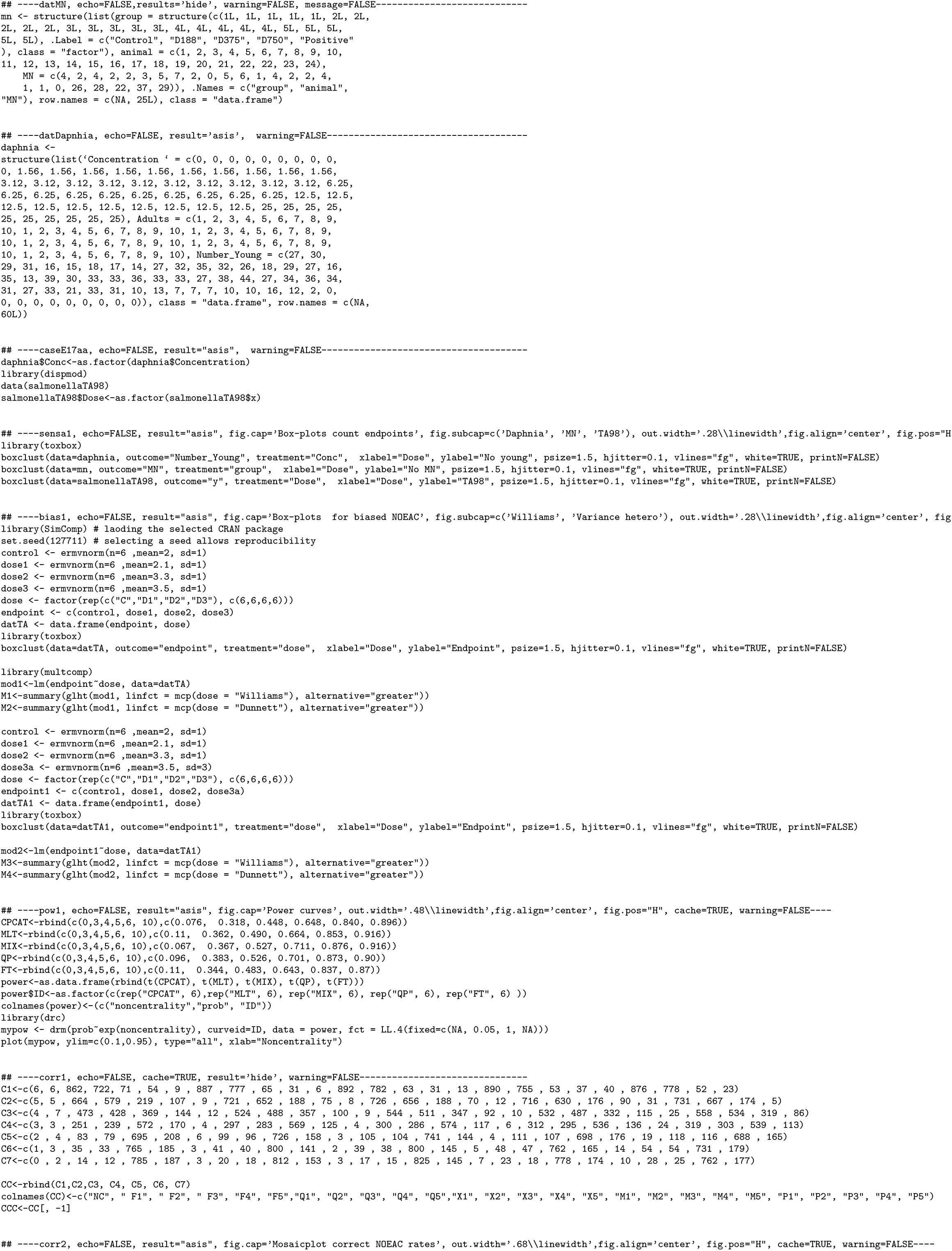
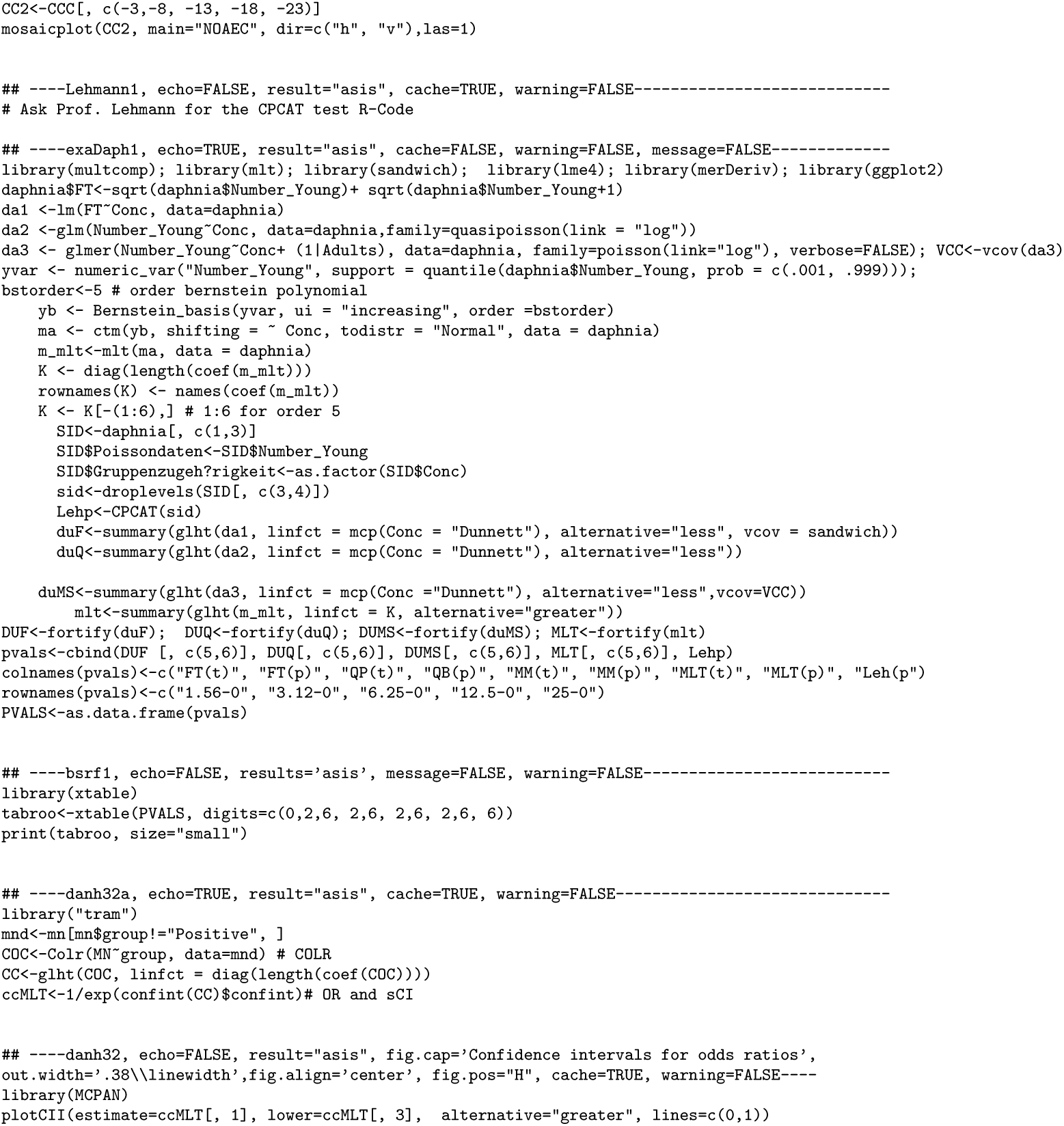

